# Accounting for individual-specific variation in habitat-selection studies: Efficient estimation of mixed-effects models using Bayesian or frequentist computation

**DOI:** 10.1101/411801

**Authors:** Stefanie Muff, Johannes Signer, John Fieberg

## Abstract

1. Popular frameworks for studying habitat selection include resource-selection functions (RSFs) and step-selection functions (SSFs) estimated using logistic and conditional logistic regression, respectively. Both frameworks compare environmental covariates associated with locations animals visit with environmental covariates at a set of locations assumed available to the animal. Conceptually, random coefficients could be used to accommodate inter-individual heterogeneity with either approach, but straightforward and efficient one-step procedures for fitting SSFs with random coefficients are currently lacking.

2. We take advantage of the fact that the conditional logistic regression model (*i. e.*, the SSF) is likelihood-equivalent to a Poisson model with stratum-specific intercepts. By interpreting the intercepts as a random effect with a large (fixed) variance, inference becomes feasible with standard Bayesian techniques, but also with frequentist methods that allow one to fix the variance of a random effect. We compare this approach to other commonly applied alternatives, including random intercept-only models, and to a two-step algorithm for fitting mixed-effects models.

3. We also reinforce the need to weight available points when fitting RSFs, since models fit using “infinitely weighted logistic regression” have been shown to be equivalent to an inhomogeneous Poisson process (IPP). We generalize this result to “infinitely weighted Poisson regression”, which converges to the same underlying IPP distribution.

4. Using data from Eurasian otters (*Lutra lutra*) and mountain goats (*Oreamnos americanus*), we illustrate that our models lead to valid and feasible inference. In addition, we conduct a simulation study to demonstrate the importance of including random slopes when estimating individual- and population-level habitat-selection parameters.

5. By providing coded examples using integrated nested Laplace approximations (INLA) and Template Model Builder (TMB) for Bayesian and frequentist analysis via the R packages `R-INLA` and `glmmTMB`, we hope to make efficient estimation of RSFs and SSFs with random effects accessible to anyone in the field. SSFs with individual-specific coefficients are particularly attractive since they can provide insights into movement and habitat-selection processes at fine-spatial and temporal scales, but these models had previously been very challenging to fit.

## 1 Introduction

Ecologists have long been interested in understanding how animals select habitat and the resulting fitness consequences from different space-use strategies. Habitat-selection analyses typically quantify preference for various resources by comparing environmental covariates at visited locations to environmental covariates at a set of locations assumed available to the animal (Manly et al., 2002). Most telemetry studies result in locations from multiple animals (*e. g.* equipped with Very High Frequency (VHF) radio collars or Global Positioning System Satellite (GPS) collars), providing an opportunity to study individual variation in habitat-selection strategies and the resulting consequences for animal fitness (Leclerc et al., 2016).

Regression models that incorporate random effects offer a powerful approach to studying inter-individual variability and are frequently used to accommodate non-independent data in ecological studies (Fieberg et al., 2009; Muff et al., 2016). Gillies et al. (2006) recommended using random intercepts to account for unequal sample sizes in habitat-selection studies, and random slope coefficients (equivalently denoted as *random coefficients* or *random slopes*) to account for individual-specific differences in habitat selection. Similarly, Hebblewhite and Merrill (2008) recommended random intercepts to account for correlation within nested groupings of locations from socially-structured populations (*e. g.* repeated observations from individual wolves and observations from wolves in the same pack). Gillies et al. (2006) and Hebblewhite and Merrill (2008) further emphasized that random coefficients could be used to model variation in habitat selection attributable to differences in habitat availability – *i. e.*, functional responses (Mysterud and Ims, 1998). Soon thereafter, Matthiopoulous et al. (2011) and Aarts et al. (2013) developed a formal framework for modelling functional responses using a combination of random effects and fixed effects constructed from the first few moments (mean, variance) of habitat covariates.

The above papers all focus on what Johnson (1980) called 3rd order selection, with available points sampled randomly or systematically from within an animal’s estimated home range. In the wildlife literature, the combined observed and available locations are typically analyzed using logistic regression, with specific focus on estimating the exponential of the linear predictor (with the intercept removed), referred to as a resource-selection function (RSF). Warton and Shepherd (2010) provided context for interpreting RSFs by showing that slope parameters in logistic regression models are asymptotically equivalent to slope parameters in an inhomogeneous Poisson point process (IPP) model. Thus, regression parameters describe relationships between environmental covariates and the relative density of observed locations in space. Fithian and Hastie (2013) further showed that equivalence between logistic regression and an IPP only holds when the model is correctly specified or when available points are “infinitely” weighted. Interestingly, several other modelling approaches, including the maximum entropy method (Maxent, Phillips et al., 2006), weighted distribution theory (Lele and Keim, 2006), and resource utilization functions (Millspaugh et al., 2006) have also been shown to be equivalent to fitting an IPP model (Aarts et al., 2012; Fithian and Hastie, 2013; Hooten et al., 2013; Renner and Warton, 2013).

Most modern statistical software platforms provide methods for fitting generalized linear mixed effects models (*e. g. logistic regression* with random intercepts and slopes), and therefore, allow for the possibility of studying individual-specific variation in studies focused on 3rd order habitat selection. However, Fieberg et al. (2010) questioned whether random intercepts are effective, or even necessary to include, when attempting to account for within-individual autocorrelation. In particular, they pointed out that the intercept depends on the probability of a location being used (relative to available), which is something under control of the analyst. We will come back to this point later in the paper.

Recent methodological development has focused on modelling habitat selection at finer temporal and spatial scales, in part driven by concerns associated with non-independence of animal locations (Arthur et al., 1996; Rhodes et al., 2005; Fortin et al., 2005). Recognizing that not all areas of the home range are equally available at all time points, Fortin et al. (2005) suggested resampling step lengths (distances between successive observed locations) and turn angles (deviations from previous bearings) to generate random movements and hence available points conditional on the previously observed location. This process results in stratified datasets with a different set of available points associated with each observed location. The combined (stratified) observed and available location data are typically analyzed using conditional logistic regression, with the exponential of the linear predictor referred to as a step-selection function (SSF). Forester et al. (2009), Duchesne et al. (2015) and Avgar et al. (2016) further refined this approach and demonstrated the utility of using common statistical distributions to model and simulate step lengths and turn angles. Specifically, they showed that it was possible to fit the equivalent of a biased random walk model when random points were generated using specific statistical distributions and when movement-related covariates (*e. g.* turn angles, step length, log-step-length) were included in conditional logistic regression models. These methods have recently been implemented in the `amt` R package (Signer, 2018; Signer et al., 2018), making SSFs an exciting and accessible approach for studying habitat selection at the scale of the movement step.

Duchesne et al. (2010) demonstrated the utility of random coefficients when modelling fine-scale habitat selection via SSFs. In particular, mixed conditional logistic regression models allow the influence of habitat covariates to depend on what is available to the animal, that is, they allow for functional responses. Unfortunately, these models are extremely challenging to fit, especially with large numbers of strata (Craiu et al., 2011). To circumvent this problem Craiu et al. (2011) developed a two-step estimation approach to fitting mixed-effects models. This approach works well when the number of strata per individual is large, but can fail (or lead to numerical instabilities) when including categorical predictors. For instance, it is not possible to use this approach in cases where one or more individuals do not encounter all factor levels of a categorical predictor.

Our objectives for this paper are to: 1) review current use of mixed effects modelling in the context of habitat-selection studies; 2) reiterate the importance of including random coefficients (not just random intercepts) in habitat-selection models; 3) reiterate the importance of weighting available points when fitting RSFs; and 4) develop computationally efficient and consistent methods for fitting RSFs and SSFs with random effects. To allow fitting of SSFs, we propose to reformulate the conditional logistic regression model as a (likelihood-equivalent) Poisson regression, where stratum-specific intercepts are modeled as a random effect with a fixed large prior variance, namely to avoid shrinkage. In addition, we explain why, for the same reason, random intercept variances in RSFs should also be fixed at a large value. We illustrate that these models are easy to fit in R (R Core Team, 2018), either employing a Bayesian approach via integrated nested Laplace approximations (INLA, Rue et al., 2009) using the R-interface `R-INLA`, or in a frequentist approach using Template Model Builder (TMB) via the `glmmTMB` R-package (Brooks et al., 2017; Magnusson et al., 2017). To illustrate the efficiency and accuracy of these methods, we reanalyzed data from a study on mountain goats (*Oreamnos americanus*) and Eurasian otters (*Lutra lutra*), and carried out a simulation study. We also compare these methods to existing two-step procedures. To facilitate uptake of these methods, we include ready-to-use R code to replicate all of our analyses.

## 2 Literature review

We reviewed the current literature to quantify the use of random effects in habitat-selection studies. We focused our review on RSFs because of the computation challenges of fitting SSFs with random effects; for many applications two-step methods offer the only feasible approach (Craiu et al., 2011). We were interested in model specification (*i. e.*, did studies use random intercepts, random slopes, or both?) and the statistical framework used for parameter estimation (Bayesian or frequentist). We downloaded all publications that cited Gillies et al. (2006) from Jan 2016 to May 2018 (*n* = 121), and then selected application papers that were peer-reviewed, written in English, fit an RSF, and claimed to use random effects (*n* = 69). Of these 69 publications, 68.1 % used a random intercept, and 18.9 % used both a random intercept and random slopes. For the remaining 13 %, it was not possible to determine from the publication text if they used random intercepts, random slopes or both, as they did not describe the model specification in further detail. With regard to the estimation method, we found a strong preference for frequentist Maximum Likelihood (ML) based estimation (72.5 %) over Bayesian methods (8.7 %). For the remaining 18.8 % of the publications it was not clear from the text which method was used for parameter estimation.

## 3 Background on analyzing RSFs and SSFs

Both RSFs and SSFs quantify habitat selection by comparing environmental covariates associated with locations that animals visit (encoded as *y* = 1) with environmental covariates at a set of locations assumed available to the animal (encoded as *y* = 0). The main difference between the RSF and the SSF approach is that the latter conditions the set of available points on the current location of the animal, resulting in a stratified dataset, whereas RSFs use a single set of (pooled) available locations for each animal, with these locations usually generated by sampling randomly or systematically from within an animal’s home range (*e. g.* Manly et al., 2002). The sampling scheme used to generate available points dictates how the respective data should be analyzed (Warton and Aarts, 2013). RSFs can be estimated by fitting a standard logistic regression model (*e. g.* Warton and Shepherd, 2010; Fithian and Hastie, 2013). On the other hand, SSFs need to account for the fact that a unique set of available points is chosen for (or”matched” to) each observed location, which can be accomplished by fitting a *conditional logistic regression* model. Each observed location thus forms a stratum along with its set of matched available locations, with exactly one point per stratum selected by the individual (McDonald et al., 2006; Duchesne et al., 2010).

### 3.1 Unmatched designs (RSFs)

Assume we have *n* = 1,…, *N* individuals and *j* = 1,…, *J*_*n*_ locations that are either used by or available to animal *n*. In the absence of any random effects, the probability that a point *y*_*nj*_ with covariate vector ***x***_*nj*_ is used, `Pr`(*y*_*nj*_ = 1 | ***x***_*nj*_) = *π*_*nj*_, can then be modeled as

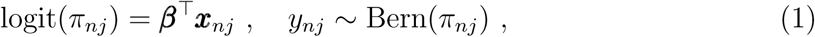

with logistic (logit) link and covariate vector ***β*** that is the target of interest (Warton and Shepherd, 2010). Standard generalized linear model (GLM) software, such as the `glm()` function in R, can be used to estimate ***β***. An extension of model (1) to include individual-specific random effects is conceptually straightforward, and the respective mixed model can for example be fit by the `glmer()` function from the `lme4` package (Bates et al., 2015).

It is important to note that, unlike prospective sampling designs involving a binary response variable, the *y*_*nj*_ in unmatched RSF designs are not Bernoulli random variables. Rather, the Bernoulli likelihood formed by (1) results in a set of estimating equations that produce consistent estimators of ***β*** in an equivalent log-linear IPP model (Warton and Shepherd, 2010). This equivalence holds whenever the RSF model is correctly specified or when available points are given infinite weights, as described by Fithian and Hastie (2013). For the respective weighted logistic regression approach, the likelihood for the available “background” samples (*i. e., y* = 0) is weighted with a weight *W*, while the used points (*y* = 1) keep weight 1. Fithian and Hastie (2013) demonstrated how, for *W* → ∞ the likelihood converges to the IPP likelihood, but in our experience values of *W* = 1000 typically lead to good approximations. If uncertainty remains, larger values may be tried for comparison. Weights are easily incorporate into most GLM software (e.g., `glm()` or `glmer()`). We do not reiterate the logistic regression likelihood here, but refer the reader to Hosmer and Lemeshow (2000) for more on logistic regression, and to Warton and Shepherd (2010) and Fithian and Hastie (2013) for a description and justification of its use for studying habitat selection.

### 3.2 Matched designs (SSFs)

In contrast to unmatched RSF designs, it seems more challenging to fit a conditional logistic regression model that emerges in the context of an SSF. Assume we have *n* = 1,…, *N* individuals with realized steps at time points *t* = 1,…, *T*_*n*_, with *j* = 1,…, *J*_*n,t*_ locations that were either used or available to animal *n* at time step *t*. Note that, for notational simplicity, we may replace *J*_*n,t*_ by *J*, because it is common practice to match a constant number of available points to each observed location. Used and available locations associated with each step form a choice set or *stratum*. This implies that the probability the *n*^th^ animal selects the *j*^th^ unit with habitat-specific covariates ***x***_*ntj*_ at time point *t* is

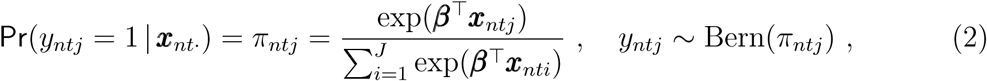

with covariate vector ***β*** that is the target of estimation. Probably the currently most popular and computationally most efficient way to fit the discrete choice model (2) in the context of habitat-selection studies is by interpreting it as a specific version of the stratified proportional hazards model (Manly et al., 2002; McDonald et al., 2006). In the absence of random effects, this “Cox trick” provides a framework for efficient inference using ML, for instance by using the `clogit()` function from the `survival` package in R (Therneau, 2015b). Unfortunately, however, the approach breaks down when random effects enter the model.

## 4 Mixed effects modelling of RSFs and SSFs

### 4.1 The importance of random slopes

Virtually all resource selection studies monitor multiple animals, and the respective data are combined and modeled jointly. However, it is well known that such a sampling design generally leads to non-independence among data points (see *e. g.* Gillies et al., 2006; Duchesne et al., 2010; Fieberg et al., 2010). In particular, resource selection will not be constant across individuals, namely due to individual-specific preferences and/or functional responses in habitat selection (Mysterud and Ims, 1998; Hebblewhite and Merrill, 2008; Matthiopoulous et al., 2011; Aarts et al., 2013; Matthiopoulos et al., 2015). Generalized linear mixed models (GLMM) offer a powerful approach that can account for correlations induced by repeated measures designs (often denoted as *pseudoreplication*), where “repeats” in resource selection studies typically denote the ensemble of measurements taken on the same animal.

Our literature review (Section 2) suggests that it is common to include individual-specific random intercepts, but not random slopes when modelling habitat selection. This is remarkable for three reasons: First and most importantly, random intercept-only models cannot (by definition) account for among-animal variation in the regression *slopes*, that is, they cannot account for functional responses, and therefore parameter estimators are likely biased. Second, omitting random slopes induces too little uncertainty in the estimated parameters (*e. g.* Schielzeth and Forstmeier, 2009), thus it is possible that researchers end up with too high confidence in their potentially biased estimators of effect sizes. The problem is particularly acute when there are lots of observations for each animal, which is typically the case in telemetry studies. And third, the intercept reflects the probability of a location being used when all covariates are set equal to 0, and is thus heavily influenced by the sampling ratio of used versus available points (Fieberg et al., 2010). Given constant ratios for all animals (which is often the case and assumed here), it is not surprising that random intercepts will sometimes return an among-animal variance component of 0. We demonstrate these issues by comparing random-intercept only and random intercept and slope models that we fit to data from mountain goats and Eurasian otters in Section 6.

### 4.2 Computational challenges

Fitting a GLMM is generally known to be a difficult and computationally demanding task, and the user can choose among various model fitting procedures. The main challenge is that random effects lead to likelihoods given by integrals without closed form solutions. Thus, maximizing the likelihood requires numerical integration techniques (e.g., quadrature-based methods) or, alternatively, the function to be integrated can be approximated so that a closed form solution exists (an overview is given by *e. g.* Bolker et al., 2009, Table I). Note, however, that while standard logistic mixed models (*i. e.*, RSFs) can be fit with several available software packages and functions (such as `lme4::glmer()`), random effects modelling is even more challenging for matched (SSF) designs, that is, for conditional logistic regression, especially when the number of cases per stratum is greater than 1, or when the strata are unbalanced (Craiu et al., 2011). In principle, these models can be interpreted as survival models with random effects (denoted as *frailty models*), for which software solutions exist (*e. g.* Therneau, 2015a; Elff, 2016), but computation quickly becomes prohibitive for telemetry data with large numbers of strata.

Given the challenges with fitting mixed conditional logistic regression models, it is not surprising that several approaches to circumvent direct random effects estimation have been proposed, such as the use of generalized estimating equations (GEEs, Craiu et al., 2008) or a two-step estimation approach (Craiu et al., 2011). GEEs, however, provide marginal parameter estimates that tend to underestimate the true effect sizes experienced by individual animals (Lee and Nelder, 2004; Fieberg et al., 2009; Muff et al., 2016); thus, we do not generally recommend them for habitat-selection studies. The two-step approach is an efficient alternative that combines estimates of individual-specific regression parameters from standard ML methods for independent data with an expectation-maximization algorithm in conjunction with conditional restricted maximum likelihood (REML). It is available via the `Ts.estim()` function from the `TwoStepCLogit` package in R (Craiu et al., 2016). This approach is an approximate method that works best when the number of strata per animal is large (Craiu et al., 2011) and when the data are not too unbalanced (*e. g.* all animals visited all levels of a categorical covariate). However, fitting SSFs with random effects in a single modelling step is currently considered to be unfeasible with standard GLM or GLMM software.

### 4.3 An efficient alternative for SSFs

We will now illustrate how relatively simple model reformulations allow one to fit mixed conditional logistic regression models in a standard GLMM. Starting for notational simplicity with the fixed effects-only model introduced in equation (2), we take advantage of the fact that a multinomial model (of which the conditional logistic regression model is a special case, see *e. g.* McCullagh and Nelder, 1989) is likelihood-equivalent to the Poisson model

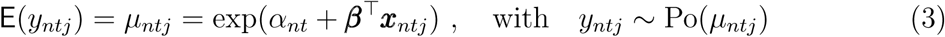

(Whitehead, 1980; McCullagh and Nelder, 1989; Baker, 1994; Chen and Kuo, 2001; Lu and Zeger, 2007), where *α*_*nt*_ is the stratum-specific intercept of animal *n* at time point *t*. Since a predefined fixed number of successes (usually one) is allowed within a stratum, the “success probability”, conditional on the outcomes of the other observations in that stratum, is thus

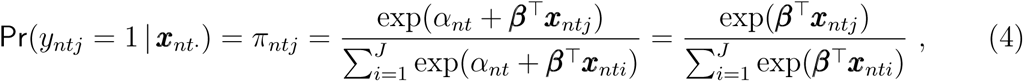

where the second equality holds because the stratum-specific intercepts *α*_*nt*_ cancel out. This illustrates that model (3) is maximizing the same likelihood-kernel as the conditional logistic model given in (2). Thus model (3), which is sometimes denoted as *conditional Poisson* model, and conditional logistic regression models give equivalent parameter estimates, 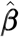, but also the same standard errors (*e. g.* McCullagh and Nelder, 1989, Chapter 6.4.2). Note that these considerations are not limited to the presence of only one used point per stratum, but are valid for any number of cases per stratum, and even hold when the different strata in a dataset contain an unequal number of cases. In addition, the reformulation also works when random effects are added to the linear predictors in (3), in which case *any* convenient GLMM software can be used to fit the resulting mixed Poisson model. This option to fit SSFs has already been pointed out by Duchesne et al. (2010), but it has only rarely been used to analyze mixed conditional logistic regression models that arise from habitat-selection studies (but see Bruun and Smith, 2003).

The obvious disadvantage of formulation (3) – and a potential reason why the approach is rarely used – is that a large number of stratum-specific intercepts *α*_*nt*_ must be estimated, which might again make the procedure prohibitive for movement data with tens of thousands of realized steps, given that each step induces a stratum. Luckily, while the *α*_*nt*_ are fixed-effects terms in the model, they are not actually of interest. It is thus possible to circumvent their explicit estimation by interpreting them as a random effect 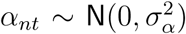, where the variance 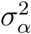 is fixed to a large value to ensure that stratum-specific intercepts are not shrunk towards their overall mean, and are thus estimated like fixed-effects parameters. This idea is easy to implement in a Bayesian approach, where such information can be specified in the priors. In fact, exactly such models have been previously implemented in a Bayesian setting under the multinomial modelling framework see *e. g.* the WinBUGS manual section 9.7 (Lunn et al., 2000). Let us finally also add random effects to the linear predictor, which leads to the mixed Poisson model

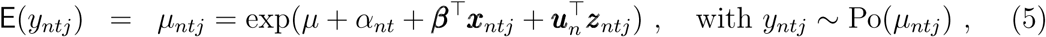

with overall mean *μ*, individual-specific random slopes 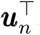, design vector ***z***_*ntj*_ (typically a sub-vector of ***x***_*ntj*_), and 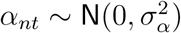 with 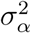 fixed at a large value, for example 10^6^. It may be prudent to verify that the results are robust when even larger values of 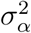 are used. Priors for the remaining parameters can be specified as in any Bayesian procedure.

Importantly, while fixing a variance in a Bayesian analysis is straightforward and natural, it is of course also possible in a likelihood framework. Model (5) can therefore also be fitted with a frequentist GLMM software, provided that there is an option to constrain 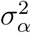 to a fixed, large value to avoid shrinkage of the intercepts. To our knowledge, this is currently not implemented in `glmer()` in the `lme4` package in R, but it is possible with the `glmmTMB` package (Brooks et al., 2017; Magnusson et al., 2017). Consequently, we will fit frequentist GLMMs using `glmmTMB::glmmTMB()`.

### 4.4 Logistic or Poisson regression?

We have now seen that a conditional logistic regression model can be reformulated as a Poisson model, which leads to feasible inference for parameters in SSFs using statistical software that allows one to fix the variance of the random intercepts. However, it may seem confusing to use a Poisson model for SSFs, while a logistic regression model is needed for RSFs. This is less surprising when considering that Poisson GLMs have been suggested for approximate inference of IPP models long before logistic regression entered the stage (Berman and Turner, 1992). Moreover and importantly, it turns out the likelihood of a Poisson model converges to the IPP likelihood when points are weighted in the manner proposed by Fithian and Hastie (2013), that is, with weights *W* → ∞ for available points and weight 1 for the observed points (as we show in Appendix 1). Consequently, RSFs can be fit using logistic *or* Poisson regression upon convenience, because the results converge to the same limit and are practically indistinguishable for large enough weights.

It may seem a logical consequence to suggest infinitely weighted Poisson regression to estimate the model parameters of equation (5) for SSFs. Unfortunately, convergence to an IPP is then not guaranteed, because the sampling of available points differs from that of the RSF, and weighting introduces an additional bias. We will illustrate this point with a simulation (see Section 6.3.2 and Figure S2 in Appendix 1), which indicates that the bias decreases for larger strata, that is, when larger numbers of available points are generated for each realized step. Here, we therefore continue with unweighted likelihoods to fit SSFs.

### 4.5 Individual-specific intercept in RSFs

As briefly mentioned in Section 4.1, the (individual-specific) intercept term in an RSF is largely determined by the sampling ratio of used and available points for each individual (Warton and Shepherd, 2010, Theorem 3.2). For example, if all covariates ***x*** in equation (1) have been mean-centered, the intercept reflects the probability that a point is used (versus available) at an “average” point in the habitat ensemble of all individuals. Thus, individual-specific intercepts may vary due to either among-individual variability in the ratio of used to available points, or due to differences in the distribution of habitat covariates within each individual’s home range (*e. g.* varying availability of woodland). Importantly, in the same way that the intercept is used to *condition on* habitat availability at the current position of an individual in an SSF, the intercept conditions on the habitat availability in the home range of the respective individual in an RSF. As a consequence, these individual-specific intercepts should *not* be shrunk towards an overall mean, but instead should also be given a large, fixed prior variance just like the stratum-specific intercepts in SSF models in Section 4.3. Alternatively, they can be treated as fixed effects (similar to other categorical covariates), which may be feasible for studies with only a few animals.

In summary, the intercepts of RSF and SSF models should both be treated in the same manner, namely as individual-specific fixed intercept effects, or equivalently, as random effects with fixed, large variance, with the latter being computationally much more efficient.

## 5 Bayesian computation for RSFs and SSFs

Our literature review suggests that in the majority of applications, RSFs and SSFs are estimated with a frequentist approach (see Section 2). Bayesian computation is thus clearly underrepresented, and it seems useful to briefly discuss some computational and conceptual aspects of the latter. Note, however, that a thorough comparison of frequentist and Bayesian paradigms is not the scope of this paper. Both have their own intrinsic advantages and disadvantages, and the choice may also depend on the preference or the background of the analyst.

The Bayesian route is often perceived to be more challenging, including reservations regarding the need to specify priors. However, thanks to the availability of Markov chain Monte Carlo (MCMC) samplers like JAGS (Plummer, 2003) or Stan (Carpenter et al., 2017), or the computationally efficient alternative provided by INLA, Bayesian computation has become increasingly accessible and more popular in the past decade. An advantage of the INLA approach used here over existing algorithms is that it circumvents time-consuming MCMC sampling by providing efficient approximations of marginal posterior distributions, and it has proven to be particularly useful for fitting GLMMs (Fong et al., 2010; Wang et al., 2018), spatial and space-time models (Blangiardo et al., 2013; Bakka et al., 2018), for modelling abundance data collected using distance sampling (Yuan et al., 2016), and for modelling species distributions more generally (Illian et al., 2013; Bakka et al., 2016). Here we discuss how to take advantage of INLA via its R inter-face `R-INLA` in the context of RSF and SSF modelling. From Sections 3 and 4 we know that weighted versions of logistic or Poisson regression models are required to estimate RSF parameters, and it is straightforward to incorporate the respective weights into the likelihood that is evaluated by `R-INLA`.

Priors for the parameters in ***β*** can be specified upon convenience. As is common, we used independent priors 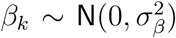 with large prior variance for all components *β*_*k*_. For the priors on the variances of the random slopes, we used penalized complexity (PC) priors. These were recently proposed as robust and intuitive alternatives to inverse gamma priors, and were shown to have excellent robustness properties (Simpson et al., 2017). PC priors are parameterized as PC(*u, α*), where the interpretation of the parameters (*u, α*) is that `Pr`(*σ* > *u*) = *α* for the standard deviation *σ*, thus the user can specify how likely it is (0 < *α* < 1) that *σ* is larger than a specific value *u* > 0. Note that we use independent priors on the random effects throughout the paper. Accounting for dependencies among the random coefficients through covariance parameters is possible, although the number of hyperparameters (variances and covariances) then grows quickly. As a consequence, the computation using INLA may become inefficient, because it is an intrinsic assumption that latent variables form a Gaussian random field, and that each non-Gaussian parameter, such as a variance, is treated as a hyperparameter over which the procedure needs to numerically integrate. Efficiency thus strongly depends the number of hyperparameters, and it should not exceed 20 (Rue et al., 2016).

## 6 Applications

### 6.1 Habitat selection of mountain goats: an RSF analysis using unmatched data

To reiterate the problems with fitting random-intercept only models, we considered data collected from GPS-collared mountain goats in British Columbia, previously analyzed by Lele and Keim (2006) and available in the `ResourceSelection` R package (Lele et al., 2017). This dataset consists of use and availability locations for each of 10 different mountain goats, with a use to available ratio of 1:2 for each goat, and a total number of 6338 used points. We first fit a RSF containing a single predictor, elevation (centered and scaled to have mean 0 and sd 1) along with a random intercept (variance not fixed) for each goat. The model was fit with an unweighted logistic regression using `glmmTMB::glmmTMB()`, and returned a variance estimate for the among-animal variability in intercepts very close to 0 (Table 1, model M1). Interestingly, a variance estimate of exactly 0 was obtained when using the `lme4::glmer()` function (results not shown).

**Table 1:** Results from four models fit to data from 10 GPS-collared mountain goats. Models M1 – M3 were fit with an unweighted likelihood. Model M4, which is the recommended model, was fit with weighted logistic regression (*W* = 1000) and fixed intercept variance 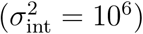. All models were fit using `glmmTMB()`.

| **Model** | $\hat{\beta}_{\text{ele}}$ | $\hat{\beta}_{\text{asp}}$ | $\hat{\sigma}_{\text{intercept}}^2$ | $\hat{\sigma}_{\text{ele}}^2$ | $\hat{\sigma}_{\text{asp}}^2$ |
| --- | --- | --- | --- | --- | --- |
| M1 (Random intercept) | 0.12 (0.05) |  | 0.008 |  |  |
| M2 (Random intercept) | 0.14 (0.03) | 0.52 (0.02) | 0.013 |  |  |
| M3 (Random intercept + slopes) | 0.07 (0.38) | 0.66 (0.11) | 0.96 | 1.40 | 0.10 |
| M4 (Random intercept + slopes) | 0.12 (0.31) | 0.65 (0.11) |  | 0.93 | 0.12 |

We next considered RSFs that included elevation plus a centered and scaled measure of aspect, and compared the estimates from a random intercept-only model (model M2) to those from a model containing both independent random intercepts and slopes (model M3), both fit with `glmmTMB()`. In the latter model, the standard errors associated with the slope coefficients for aspect and elevation were an order of magnitude larger when they were modeled as random effects. These results clearly demonstrate the problems noted by Schielzeth and Forstmeier (2009), namely that random intercept-only models tend to underestimate standard errors of (potentially biased) fixed effects parameters. Finally, we fitted the weighted logistic regression model (using *W* = 1000) with random intercept and slopes, with fixed intercept variance at 10^6^ (model M4), because this is the procedure we recommend. Weighting the likelihood and fixing the variance of the intercepts led to a noticeable increase in the estimate of *β*_ele_ and a decrease in the estimate of 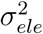, while it had little effect on the estimated values of *β*_asp_ and 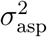. This example illustrates how an inappropriate analysis can lead to misleading conclusions, but that fitting the right model requires only little additional effort.

### 6.2 Habitat selection of otters: an SSF analysis using matched data

We reanalyzed data collected and presented by Weinberger et al. (2016) involving nine radio-collared otters that were tracked between six months and three years in the European Alps. To fit SSFs to these data, each observed location was matched with nine random (available) points generated by resampling step lengths and turning angles from their empirical distribution (Fortin et al., 2005). This process naturally resulted in a matched sampling design, with each step inducing a stratum consisting of 10 (1 used and 9 available) locations, thus a conditional regression model was needed. Due to the absence of an efficient alternative, the original analysis was performed with a two-step estimation method provided by the `TwoStepCLogit::Ts.estim()` function. The original model included 12 covariates and random effects for all of them. Here, however, we only included the variables of main interest, namely the factorial covariate *habitat type* (with levels *main discharge, reservoir* and *residual water*), the continuous variable *river width*, plus a variable that accounts for step length to reduce bias, as suggested by Forester et al. (2009). The data contained a total of 41 670 data points with 4 167 realized steps, where the latter thus corresponds to the number of strata.

For illustration, we started by fitting fixed effects-only models. To this end, the well established stratified Cox model was fit via the `survival::clogit()` function. The respective results were compared to the outcome from the conditional Poisson model as given by equation (3), where the stratum-specific intercepts are implicitly estimated by modelling them as a random intercept with a fixed variance *α*_*nt*_ *∼* `N`(0, 10^6^). We estimated the parameters both with the frequentist approach using `glmmTMB`, and with the Bayesian approach using `R-INLA`, with independent ***β*** *∼* `N`(0, 10^4^) priors for all components in the vector of slope parameters. Importantly, this led to parameter estimates that were essentially indistinguishable from those obtained via the stratified Cox model (Table 2), confirming that the conditional Poisson model is equivalent to the conditional logistic model, and that we can circumvent the explicit estimation of the stratum-specific intercepts (denoted as *α*_*nt*_ in equation 3) by a random effect with large fixed variance. Note that this equivalence does not hold when 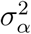 is freely estimated instead, and that this would lead to invalid results, as will be illustrated in the simulation below (Section 6.3.2). Computation times were on the order of a few seconds for all procedures.

**Table 2:** Estimated fixed effects and variance parameters of the Eurasian otter example when using the Cox model (clogit), the Poisson model with stratum-specific intercept (cPois) fit with `R-INLA` or `glmmTMB()`, and the two-step procedure `Ts.estim()` (Two-step). For the INLA output, posterior means are given for the slope estimates, and posterior modes for the variances. Values in brackets are standard errors (for the slope estimates) and 95% credible intervals (for the variances); `Ts.estim()` and `glmmTMB()` do not provide measures of uncertainty for variance parameters.

| **Slope estimates** | $\beta_{\text{STAU}}$ | $\beta_{\text{REST}}$ | $\beta_{\text{Width}}$ |
| --- | --- | --- | --- |
| **I. Fixed effects models** |  |  |  |
| clogit | −0.07 (0.07) | −0.38 (0.10) | 0.16 (0.04) |
| cPois (INLA) | −0.07 (0.07) | −0.38 (0.10) | 0.16 (0.04) |
| cPois (glmmTMB) | −0.07 (0.07) | −0.38 (0.10) | 0.16 (0.04) |
| **II. Mixed effects models (random intercept & slopes)** |  |  |  |
| Two-step | 0.04 (0.17) | −0.24 (0.24) | 0.10 (0.12) |
| cPois (INLA) | 0.02 (0.18) | −0.33 (0.22) | 0.11 (0.14) |
| cPois (glmmTMB) | −0.004 (0.14) | −0.35 (0.16) | 0.12 (0.11) |
| **Variance estimates**<br>(Mixed models only) | $\sigma_{\text{STAU}}^2$ | $\sigma_{\text{REST}}^2$ | $\sigma_{\text{Width}}^2$ |
| Two-step | 0.17 | 0.35 | 0.08 |
| cPois (INLA) | 0.08 (0.02,0.78) | 0.10 (0.03,1.03) | 0.05 (0.02,0.47) |
| cPois (glmmTMB) | 0.07 | 0.10 | 0.07 |

Next, we included independent individual-specific random slopes for all covariates (except for step length). We again estimated parameters with `glmmTMB` and `R-INLA`, using the conditional Poisson model (5). For the Bayesian model, the same priors as above were used, and PC(3, 0.05) priors on the precisions of the random slopes (but results were insensitive to this choice). These results were compared to the outcome of the two-step procedure via `Ts.estim()`, where it was also assumed that the random effects are independent. Note that the point estimates reported from the Bayesian models are posterior means (for the fixed effects) and posterior modes (for the variances), respectively, following the recommendations by He and Hodges (2008).

The results (Table 2) illustrate two important points: First, the inclusion of individual-specific random slopes in the Poisson regression model leads to different parameter estimates and to much larger standard errors for the fixed effects than when fixed effects-only models are used, which again confirms that fixed effects-only models might lead to invalid conclusions in the presence of inter-individual heterogeneity. And second, the reformulation of the conditional logistic regression model as a Poisson model with random stratum-specific intercept, as given in (5), leads to feasible estimation of mixed effects parameters in a single modelling step. While computations with other single-step R procedures, such as adding random effects (frailties) to survival models using `coxme::coxme()`, were un-feasible even when only 1 000 out of the more than 4 000 strata were used (we interrupted the sessions after 24h of non-convergence), `glmmTMB()` terminated in roughly 5 seconds and `R-INLA` in 90 seconds on an Intel Core i7-6500U 4 × 2.50GHz processor for the full dataset. On the other hand the `Ts.estim()` procedure was still considerably faster (on the order of 1.5 seconds), although the parameter estimates from the Poisson model are not in perfect agreement with those from the approximate two-step procedure, especially for *β*_REST_ and 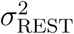.

### 6.3 Simulated data

To provide a systematic overview of the performance of the different estimation methods and model specifications, we simulated and analyzed scenarios for matched and unmatched designs with known true coefficient values. All simulations of movement tracks involved two continuous covariates: *elevation* and *habitat*. We simulated elevation and habitat as independent unconditional Gaussian Random Fields (GRF; as implemented in Ribeiro Jr and Diggle, 2016) with range *σ*^2^ = 0.1 and a partial sill of *φ* = 50. We sampled and analyzed movement data for unmatched (RSF) and matched (SSF) designs, where each setup was replicated 500 times to obtain a sampling distribution of the estimated coefficients and to investigate bias and variance of the different estimators.

#### 6.3.1 Unmatched (RSF)

We simulated data according to an unmatched sampling design by drawing used points at locations with covariate values ***x***, where animal *n* was present in the landscape with probability proportional to

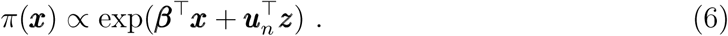

Selection coefficients were set to ***β***^*T*^ = (*β*_*ele*_, *β*_*hab*_) = (*-*4, 4) for elevation (*ele*), and habitat (*hab*), respectively. Both variables were given individual-specific slopes ***u***_*n*_, generated from uncorrelated Gaussian distributions with mean zero and variances 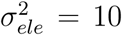 and 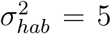. Data were generated for 20 animals and 200 locations per animal, plus nine times more (thus 1 800 per animal) background points from the landscape with uniform probability.

The simulated data were first analyzed with the correct, data-generating model that included fixed effects and independent random slopes for both covariates. For each simulated dataset, we fit mixed logistic regression models, using both unweighted and weighted likelihoods (with *W* = 1000). To illustrate that the logistic and Poisson regression models converge to the same limit for large *W*, all models were also fit using a Poisson likelihood. And finally, to illustrate how model misspecification may result in biased parameter estimators, we used weighted logistic regression to compare the fixed-effects model and a random-intercept-only model to the correctly specified mixed model. Simulated datasets were analyzed in a Bayesian framework using `R-INLA`, and by a likelihood approach using `glmmTMB`, where the individual-specific intercept variance was fixed at 10^6^ for the reason discussed in Section 4.5. For comparison, we also repeated the analyses with a freely estimated intercept variance. For the Bayesian setup, we used `N`(0, 10^3^) priors on the fixed effects, and relatively vague PC(10, 0.01) priors on 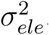, and PC(5, 0.01) priors on 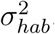.

Weighted regression models with appropriate specification of random effects led to consistent estimators, both for the logistic and Poisson likelihood, while the unweighted models resulted in biased estimators when the Poisson model was used (Figure 1, left panel). Note that the unweighted logistic model would also be expected to produce biased estimators in the presence of model misspecification (Fithian and Hastie, 2013), but given that the fitted model matches the sampling scheme (6), the unweighted logistic regression model (*i. e., W* = 1) is also consistent here. Estimators of all fixed-effects coefficients were biased in the fixed-effects-only and random-intercept-only models (Figure 2). Lastly, we obtained similar results when we treated the variance of the random intercepts as a free parameter (Figure S1 in Appendix 1). Allowing the variance of the intercepts to be estimated may introduce bias, however, for data sets with smaller (or more variable) sample sizes of observed locations and will be particularly problematic for SSFs as shown in the next section.

**Figure 1:**
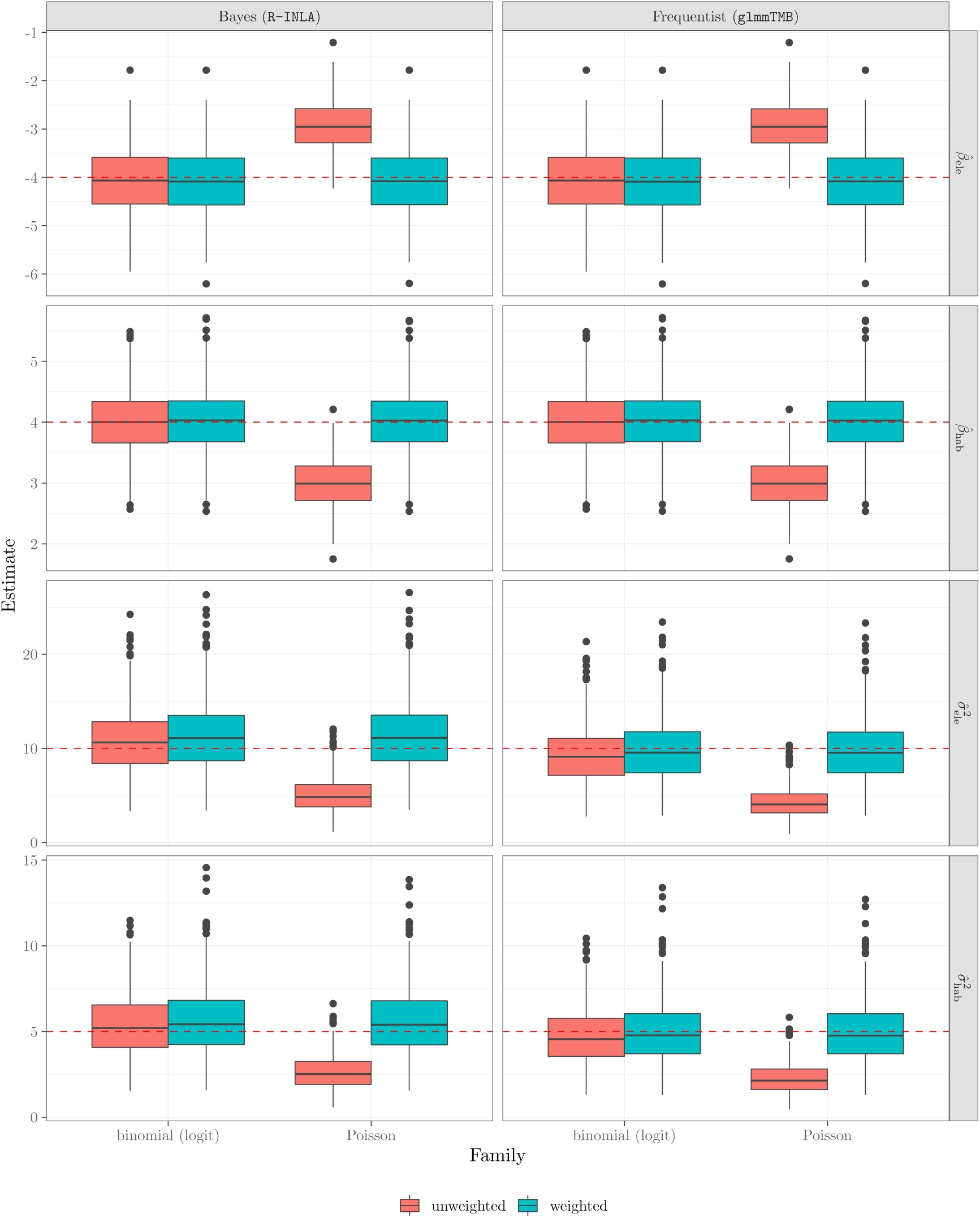
Sampling distribution of estimated RSF coefficients from the model that correctly included random intercept and random slopes, for weighted and unweighted versions (*W* = 1000) and either using a Bayesian (left) or ML approach (right). The variance of the random effect for the intercept was fixed at 10^6^. Boxplots show the distribution of 500 replications, where posterior means were used for the fixed effects and posterior modes for the variances in the Bayesian case. The horizontal red dashed lines indicate the true value used to generate the data in the simulations.

**Figure 2:**
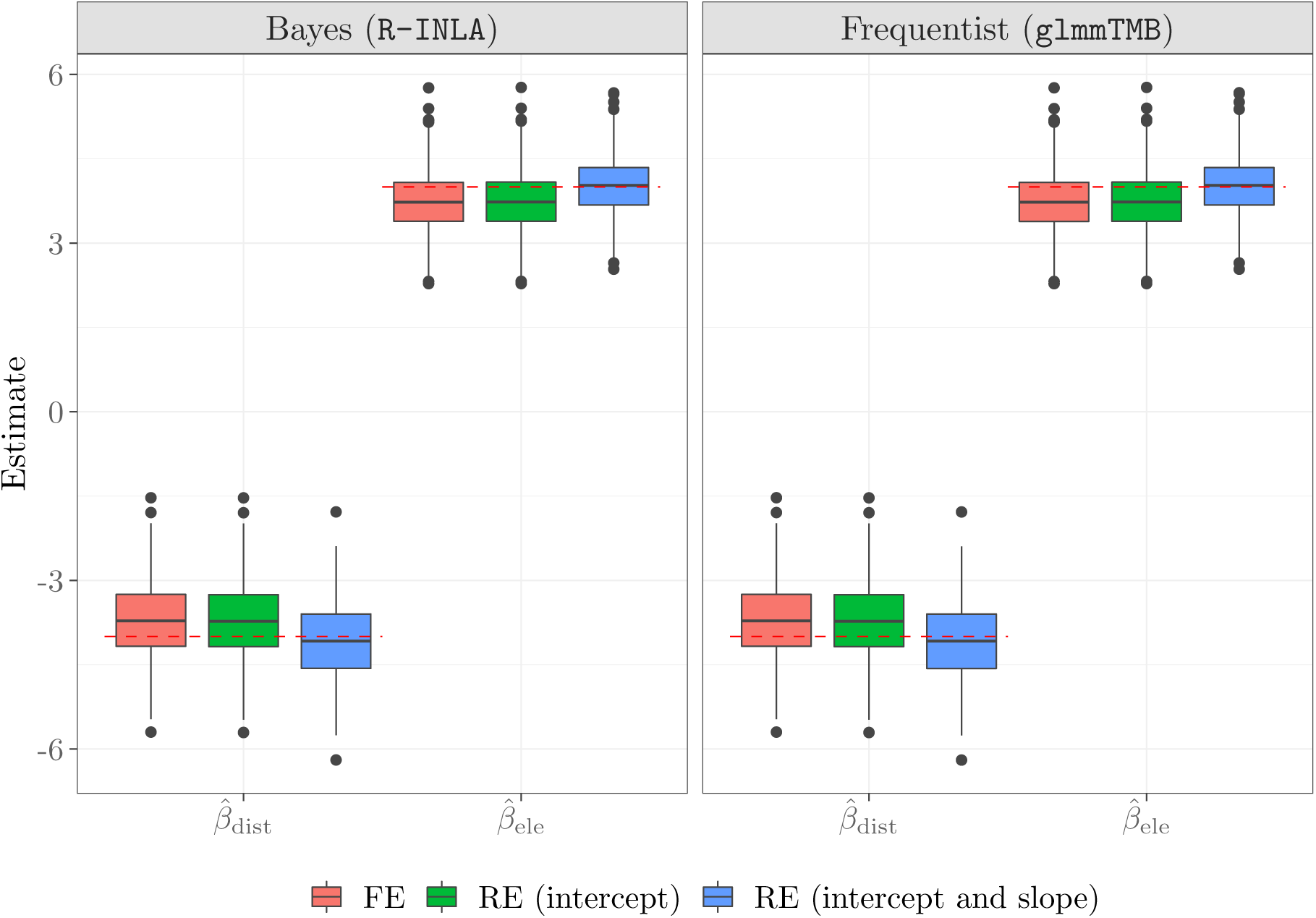
Sampling distribution for estimated RSF coefficients from weighted logistic regression models including only fixed effects (FE), or including also random effects (RE), either only a random intercept, or both a random intercept and random slopes. Shown are only models with weights (*W* = 1000) and fixed intercept variance of 10^6^ (for random intercept and random slope models only) estimated with `R-INLA` and `glmmTMB`. In the Bayesian case, the distribution of posterior means is shown. Boxplots show the distribution of 500 replications. The horizontal red dashed lines indicate the true value used to generate the data in the simulations.

These results confirm and highlight: 1) the importance of using weighted likelihoods, 2) that both weighted logistic and weighted Poisson regression can be used to analyze unmatched RSF data, and 3) the importance of including cluster-specific random slopes in real applications, given that animals living in different landscapes will usually exhibit different habitat-selection patterns.

#### 6.3.2 Matched (SSF)

To compare different estimation approaches for SSFs, that is, for matched sampling designs, we simulated movements of 20 animals according to a biased random walk starting at the center of the landscape at time *t* = 0. To find the position at time *t* + 1, each animal *n* was given 200 candidate locations, where the coordinates for each candidate location were determined by drawing a random step length from an exponential distribution with rate parameter *λ* = 1, and a random turning-angle from a uniform distribution. One candidate location was then selected at random with probability proportional to exp(***β***^*T*^***x***), where ***x*** are the covariate values at the end point of each potential step and ***β***^*T*^ = (*-*4, 4) was again the vector of selection coefficients. Animals were again assigned individual-specific slopes for both variables, generated from uncorrelated Gaussian distributions with mean ***β*** and variances 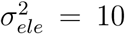 and 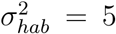. For each animal, we simulated 200 time steps, and each observed step was paired with 9 random (control) steps. Following Forester et al. (2009), we generated random steps with step lengths from an exponential distribution with rate 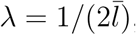 with 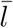 equal to the mean realized step length, and with the direction of random steps drawn from a uniform distribution distribution of turning angles between *-π* and *π*. We then included step length (*l*) in the linear predictor to correct for the bias due to the way we generated random step lengths (*i. e.*, exponential with 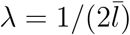 rather than *λ* = 1).

These data were analyzed with the mixed conditional Poisson model of equation (5) using frequentist (`glmmTMB`) and Bayesian (`R-INLA`) inference including random slopes for elevation and habitat. The variance of the stratum-specific intercept was fixed to 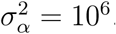. To illustrate that fixing this variance is important, we also fit the same model with 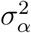 estimated instead (only with `glmmTMB` to avoid redundancy). For INLA we again used `N`(0, 10^3^) priors on the fixed effects, and 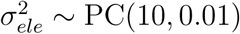 and 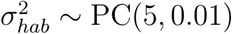 priors on the random effects. As a comparison, we also estimated regression parameters using the two-step approach implemented in `Ts.estim()` assuming independent slopes, and fit fixed-effects models with Cox models using the `clogit()` function. All these models were fit with unweighted likelihoods for the reason mentioned in Section 4.4.

As before, the Poisson models with fixed 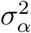 fit with `glmmTMB` and `R-INLA` retrieved consistent estimators of the fixed-effects parameters, and the two-step estimator was also nearly unbiased (Figure 3). This was not true, however, when the stratum-specific intercept variance was estimated by the Poisson model rather than fixed, in which case all estimators were heavily biased. Importantly, we again observe that ignoring random effects leads to biased estimators of fixed-effects parameters when there is inter-individual heterogeneity in the slopes. All variance estimators were slightly underestimated for all methods, namely because the step-length variable in the predictor absorbs some of the variability in the selection coefficients. Weighted regression models also resulted in biased estimators except for very large numbers of random steps (Figure S2 in Appendix 1); therefore, weighted alternatives were not further investigated here.

**Figure 3:**
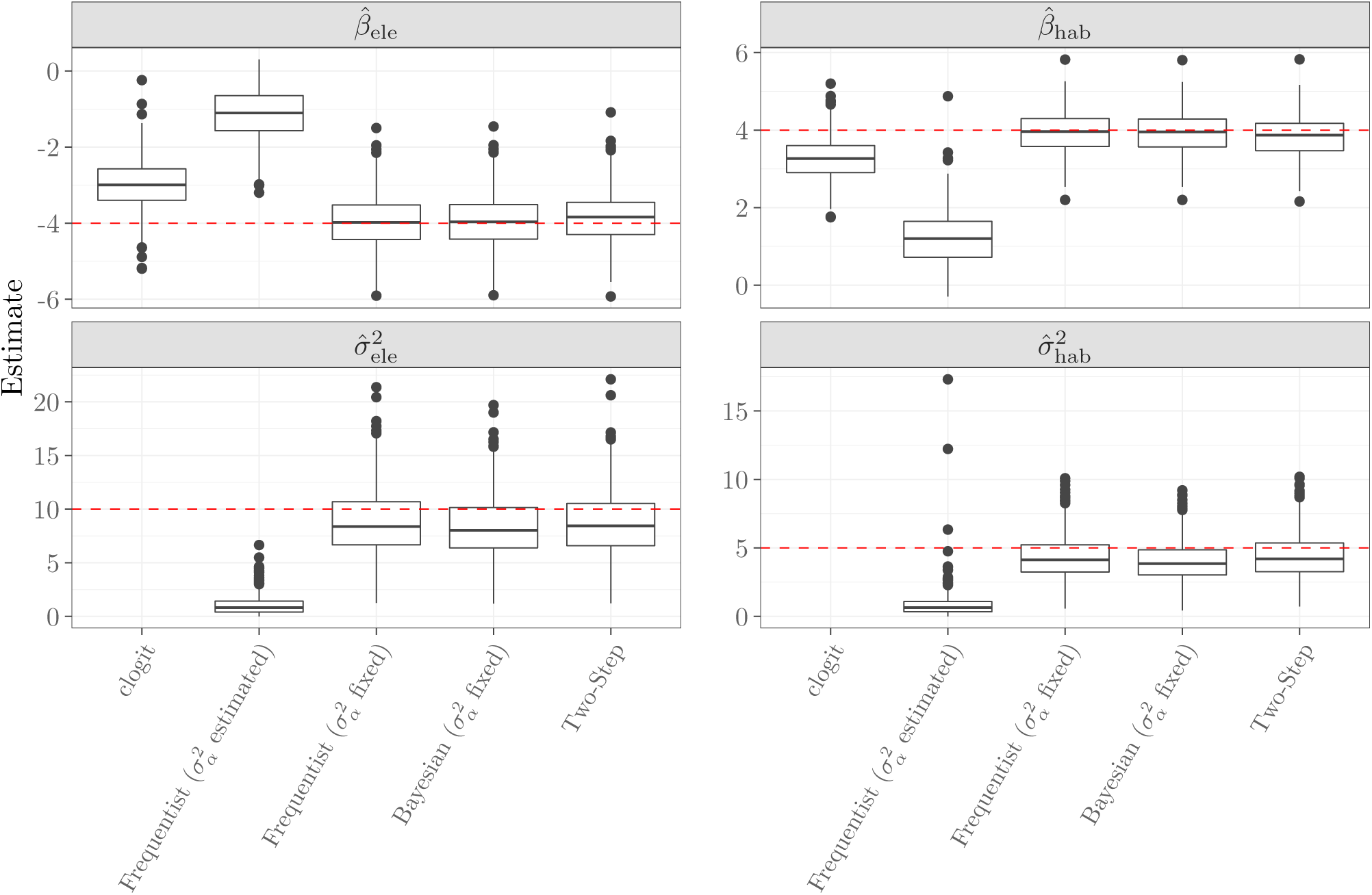
Sampling distribution for estimated SSF coefficients from the conditional Poisson regression model using a frequentist approach (`glmmTMB`), a Bayesian approach (`R-INLA`), the two-step approach implemented in the `Ts.estim()` function, and conditional logistic regression (clogit) without random effects. In the Bayesian case, posterior means are shown for the fixed effects and posterior modes for the variances. The frequentist approach was implemented both with 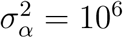 fixed (as recommended) or by estimating 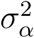 (for illustration). Boxplots show the distribution of 500 replications. The horizontal red dashed lines indicated the true value used for the simulations.

## 7 Discussion

Recent technological advances have made it possible track a wider range of species for longer durations, leading to an explosion of high-temporal resolution location data (Kays et al., 2015). For example, Movebank, an online platform for storing, managing, and sharing data now includes more than 700 million locations from over 4000 studies of close to 800 different taxa (Kranstauber et al., 2011; Wikelski and Kays, 2018). The widespread availability of fine-scale temporal data is fueling the development of new statistical approaches for modelling animal movement data (*e. g.* Hooten et al., 2017; Jonsen et al., 2018) and also provides unique opportunities to study among-individual variability in movement and habitat-selection patterns.

Step-selection functions were developed to address concerns regarding statistical independence in habitat selection studies that utilize fine-scale location data, and they are appealing because they provide an objective approach to determining habitat availability based on movement characteristics of the study species (Fortin et al., 2005; Thurfjell et al., 2014). Although fitting step-selection models to individual animals is straightforward, efficient estimation procedures for models fit to multiple animals have been lacking, hindering our ability to quantify among-animal variability in their habitat-selection patterns. Mixed-effects models are an attractive option, but these models are well acknowledged to be computationally challenging to fit in this context (Duchesne et al., 2010).

We proposed to fit RSFs and SSFs in a unified, standard GLMM framework, which is possible by combining three statistical results. First, we make use of the fact that the conditional logistic regression model, which needs to be fit to derive SSFs, is actually a multinomial model, and as such it is likelihood-equivalent to a Poisson model. This renders mixed-effects modelling for SSFs equivalent to fitting any Poisson GLMM, which implies that incorporating individual-specific variation in SSFs is no more challenging than doing so for RSFs. Second, because individual- or stratum-specific intercepts are not actually of interest in RSFs or SSFs, and because they are determined by sampling ratios and habitat availability, these intercepts should be treated as fixed effects, or equivalently and more efficiently, as random effects with large, fixed variance. Doing so prevents these parameters from being shrunk towards the overall mean. By contrast, treating the intercepts as random effects with estimated variance results in biased estimators of the slope parameters that are the target of inference. The magnitude of the shrinkage, and hence bias, may be minimal for RSFs that include many observations for each individual (as in the goat example of Section 6.1 and our simulation study in 6.3.2), but can be substantial for SSFs which tend to include only a few observations in each stratum (Figure 3). And third, we reiterated that the weighted logistic regression likelihood, with weights *W* on the available points, converges to the IPP likelihood for *W* → ∞ (Fithian and Hastie, 2013), and have expanded this result to the Poisson likelihood Appendix 1. RSFs can therefore both be fit with weighted logistic *or* weighted Poisson regression upon convenience.

Poisson and/or logistic regression mixed-effects models with fixed intercept variance led to consistent inference for all model parameters, both for RSFs and SSFs. Thus, we have demonstrated that it is feasible to fit SSFs with random coefficients in a single step using standard statistical software, both for Bayesian or frequentist inference. It is particularly straightforward to fix the individual- or stratum-specific intercept variance in a Bayesian framework, where the user is required to specify priors on all unknown parameters. To ensure efficient Bayesian inference we have relied on the INLA approach via the `R-INLA` interface, which in preliminary tests using simulated datasets was considerably faster than implementations using MCMC sampling via Stan (Carpenter et al., 2017). Users that prefer frequentist inference should choose a software package that allows to fix a random effect variance to a prespecified value. Here we fitted these models using `glmmTMB`, which provides fast inference, and has previously proven useful for analyzing large telemetry data sets (Jonsen et al., 2018). Table 3 gives an overview of models and procedures that we recommend for efficient and accurate inference on either fixed-effects or random-effects RSFs and SSFs.

**Table 3:** Overview of sampling designs and procedures in R that we recommend for efficient computation. Please note that we recommend to carry out RSF analyses using the *infinitely weighted* version. ^(*\**)^Poisson regression for RSFs is equivalent to logistic regression in the limit of *W* → ∞.

|  |  | Unmatched designs (RSFs) | Matched designs (SSFs) |
| --- | --- | --- | --- |
| Fixed effects | Example | Mountain goats (sec. 6.1) | Eurasian otters (sec. 6.2) |
|  | Models: | Logistic regression<br>Poisson regression<sup>(*)</sup> | Conditional logistic regression<br>Conditional Poisson regression<br>(models (2) and (3) in text) |
| | R procedures: | `inla()`, `glm()`, `glmmTMB()` | `clogit()` function<br>*or* `inla()/glmmTMB()` for Poisson models<br>with stratum-specific random effect<br>and large fixed variance $\sigma_\alpha^2$. |
| Mixed effects | Models: | Mixed logistic regression<br>Mixed Poisson regression<sup>(*)</sup> | Mixed conditional Poisson<br>regression (model (5) in text) |
|  | R procedures: | `inla()`, `glmer()`, `glmmTMB()` | `inla()`, `glmmTMB()`, `Ts.estim()` |

Prior to now, fitting random coefficient SSFs was often only feasible via two-step procedures (Craiu et al., 2011; Hooten et al., 2016). An advantage of using `Ts.estim()` is that it is typically much faster than `glmmTMB()` and (even more so) than `R-INLA`, as illustrated by the computation times of the otter data analysis of Section 6.2. As a second benchmark, we also fit an SSF model including 14 fixed and 14 random effects on a real dataset from 13 GPS-collared *Eurasian lynx* (Gehr et al., 2017) with 18 762 realized steps (strata) and a total of 144 810 data points on our Intel Core i7-6500U 4 × 2.50GHz processor with 16GB RAM. The respective `R-INLA` procedure terminated in a bit less than 10 hours, whereas the same model could be fit in 13 minutes with `glmmTMB()` and in a few seconds with `Ts.estim()`. However, it must be kept in mind that `Ts.estim()` is an approximate procedure and may fail to converge, for example, when at least one animal does not encounter and use all habitat types available. Still, for very large datasets and models, where GLMMs may demand too much computational power, it certainly remains a convenient and efficient alternative.

We have seen that frequentist analyses with `glmmTMB` can be considerably faster than the Bayesian route using `R-INLA`. In fact, efficiency gain will rarely be the reason to choose Bayesian over likelihood inference. An interesting benefit of Bayesian procedures is that they give (marginal) posterior distributions of all parameters, whereas frequentist approaches only return point estimates and standard errors for fixed effect parameters, but no measures of uncertainty for variance parameters. In addition, various modelling extensions, such as spatial or temporal dependencies (*e. g.* Lindgren et al., 2011; Blangiardo et al., 2013) or measurement error in covariates (*e. g.* Muff et al., 2015) are often much more straightforward to incorporate, or even only computationally feasible, in a Bayesian setup. For further comparison of frequentist and Bayesian approaches, we refer the reader to an extensive body of literature (*e. g.* Efron, 1986; Bayarri and Berger, 2004; Gelman et al., 2009, just to mention a few).

Although the importance of including random coefficients in regression models of habitat-selection studies has been stressed repeatedly (Gillies et al., 2006; Duchesne et al., 2010), our literature review suggests that random-effects models are often understood as models that merely include a random intercept. Here we have reiterated and illustrated that such practice may lead to too high confidence in results that are potentially biased. By providing coded examples using `R-INLA` and `glmmTMB`, we hope to make efficient estimation of RSFs and SSFs with random effects accessible to anyone in the field. SSFs with individual-specific coefficients are particularly attractive since they can provide insights into movement and habitat-selection processes at fine-spatial and temporal scales (Avgar et al., 2016; Signer et al., 2018), but these models had previously been very challenging to fit.

## Acknowledgements

JF received partial support from the Minnesota Agricultural Experimental Station and the McKnight Foundation and SM was supported by the Faculty of Science of the University of Zurich. The authors would like to thank Irene Weinberger for the permission to use the Eurasian otter dataset, and Benedikt Gehr for the Eurasian lynx data and comments on an earlier version of the manuscript.

## Author’s contributions

JF, SM and JS conceived the research idea, SM developed the statistical framework, SM, JF and JS conceived of the design and analysis of the data, JS developed and ran the simulations, JS conducted the literature review, SM and JF led the writing of the manuscript. All authors contributed critically to the drafts and gave final approval for publication.

## Data accessibility

Data and code for all examples and simulations presented here will be archived with the Data Repository of the University of Minnesota (https://www.lib.umn.edu/datamanagement/drum) upon acceptance of the paper.

## References

Aarts, G., J. Fieberg, S. Brasseur, and J. Matthiopoulos (2013). Quantifying the effect of habitat availability on species distributions. Journal of Animal Ecology 82, 1135–1145.

Aarts, G., J. Fieberg, and J. Matthipoulos (2012). Comparative interpretation of count, presence–absence and point methods for species distribution models. Methods in Ecology and Evolution 3, 177–187.

Arthur, S. M., B. F. Manly, L. L. McDonald, and G. W. Garner (1996). Assessing habitat selection when availability changes. Ecology 77, 215–227.

Avgar, T., J. R. Potts, M. A. Lewis, and M. S. Boyce (2016). Integrated step selection analysis: bridging the gap between resource selection and animal movement. Methods in Ecology and Evolution 7, 619–630.

Baker, S. G. (1994). The multinomial-Poisson transformation. Journal of the Royal Statistical Society. Series D (The Statistician) 43, 495–504.

Bakka, H., H. Rue, G. Fuglstad, A. Riebler, D. Bolin, J. Illian, E. Krainski, D. Simpson, and F. Lindgren (2018). Spatial modeling with R-INLA: A review. Wiley Interdisciplinary Reviews: Computational Statistics, e1443.

Bakka, H., J. Vanhatalo, J. Illian, D. Simpson, and H. Rue (2016). Accounting for physical barriers in species distribution modeling with non-stationary spatial random effects. ArXiv e-prints. arXiv:1608.03787v1.

Bates, D., M. Mächler, B. Bolker, and S. Walker (2015). Fitting linear mixed-effects models using lme4. Journal of Statistical Software 67, 1–48.

Bayarri, M. and J. Berger (2004). The interplay of bayesian and frequentist analysis. Statistical Science 19, 58–80.

Berman, M. and T. R. Turner (1992). Approximating point process likelihoods with GLM. Journal of Applied Statistics 41, 31–38.

Blangiardo, M., M. Cameletti, G. Baio, and H. Rue (2013). Spatial and spatio-temporal models with R-INLA. Spatial and Spatio-temporal Epidemiology 4, 33–49.

Bolker, B. M., M. E. Brooks, C. J. Clark, S. W. Geange, J. R. Poulsen, M. H. H. Stevens, and J.-S. S. White (2009). Generalized linear mixed models: a practical guide for ecology and evolution. Trends in Ecology and Evolution 24, 127–135.

Brooks, M. E., K. Kristensen, K. J. van Benthem, A. Magnusson, C. W. Berg, A. Nielsen, H. J. Skaug, M. Maechler, and B. M. Bolker (2017). Modeling zero-inflated count data with glmmTMB. bioRxiv preprint bioRxiv:132753.

Bruun, M. and H. G. Smith (2003). Landscape composition affects habitat use and foraging flight distances in breeding european starlings. Biological Conservation 114, 179–187.

Carpenter, B., A. Gelman, M. Hoffman, D. Lee, B. Goodrich, M. Betancourt, M. Brubaker, J. Guo, P. Li, and A. Riddell (2017). Stan: A probabilistic programming language. Journal of Statistical Software, Articles 76, 1–32.

Chen, Z. and L. Kuo (2001). A note on the estimation of the multinomial logit model with random effects. The American Statistician 55, 89–95.

Craiu, R. V., T. Duchesne, and Fortin (2008). Inference methods for the conditional logistic regression model with longitudinal data. Biometrical Journal 50, 97–109.

Craiu, R. V., T. Duchesne, D. Fortin, and S. Baillargeon (2011). Conditional logistic regression with longitudinal follow-up and individual-level random coefficients: A stable and efficient two-step estimation method. Journal of Computational and Graphical Statistics 20, 767–784.

Craiu, R. V., T. Duchesne, D. Fortin, and S. Baillargeon (2016). TwoStepCLogit: Conditional Logistic Regression: A Two-Step Estimation Method. R package version 1.2.5.

Duchesne, T., D. Fortin, and N. Courbin (2010). Mixed conditional logistic regression for habitat selection. Journal of Animal Ecology 79, 548–555.

Duchesne, T., D. Fortin, and L.-P. Rivest (2015). Equivalence between step selection functions and biased correlated random walks for statistical inference on animal movement. PLOS ONE 10, e0122947.

Efron, B. (1986). Why isn’t everyone a Bayesian? The American Statistician 40, 1–5.

Elff, M. (2016). mclogit: Mixed Conditional Logit Models. R package version 0.4.4.

Fieberg, J., J. Matthiopoulos, M. Hebblewhite, M. S. Boyce, and J. L. Frair (2010). Correlation and studies of habitat selection: problem, red herring or opportuniy? Phil. Trans. R. Soc. B 365, 2233–2244.

Fieberg, J., R. H. Rieger, M. C. Zicus, and J. S. Schildcrout (2009). Regression modelling of correlated data in ecology: subject-specific and population averaged response patterns. Journal of Applied Ecology 46, 1018–1025.

Fithian, W. and T. Hastie (2013). Finite-sample equivalence in statistical models for presence-only data. Annals of Applied Statistics 7, 1917–1939.

Fong, Y., H. Rue, and J. Wakefield (2010). Bayesian inference for generalized linear mixed models. Biostatistics 11, 397–412.

Forester, J. D., H. K. Im, and P. J. Rathouz (2009). Accounting for animal movement in estimation of resource selection functions: sampling and data analysis. Ecology 90, 3554–3565.

Fortin, D., H. Beyer, M. Boyce, and D. Smith (2005). Wolves influence elk movements: behavior shapes a trophic cascade in yellowstone national park. Ecology 86, 1320–1330.

Gehr, B., E. Hofer, S. Muff, A. Ryser, E. Vimercati, K. Vogt, and L. F. Keller (2017). Spatial scale and behavioral state interact in shaping temporal dynamics of habitat selection in Eurasian lynx. Oikos 126, 1389–1399.

Gelman, A., J. B. Carlin, H. S. Stern, and D. B. Rubin (2009). Bayesian Data Analysis (2 ed.). Boca Ration: Chapman & Hall/CRC.

Gillies, C. S., M. Hebblewhite, S. E. Nielsen, M. A. Krawchuk, C. L. Aldridge, J. L. Frair, D. J. Saher, C. E. Stevens, and C. L. Jerde (2006). Application of random effects to the study of resource selection by animals. Journal of Animal Ecology 75, 887–898.

He, Y. and J. S. Hodges (2008). Point estimates for variance-structure parameters in bayesian analysis of hierarchical models. Computational Statistics & Data Analysis 52, 2560–2577.

Hebblewhite, M. and E. Merrill (2008). Modelling wildlife–human relationships for social species with mixed-effects resource selection models. Journal of applied ecology 45, 834–844.

Hooten, M. B., F. E. Buderman, B. M. Brost, E. M. Hanks, and J. S. Ivan (2016). Hierarchical animal movement models for population-level inference. Environmetrics 27, 322–333.

Hooten, M. B., E. M. Hanks, D. S. Johnson, and M. W. Alldredge (2013). Reconciling resource utilization and resource selection functions. Journal of Animal Ecology 82, 1146–1154.

Hooten, M. B., D. S. Johnson, B. T. McClintock, and J. M. Morales (2017). Animal movement: statistical models for telemetry data. Boca Raton: CRC Press.

Hosmer, D. W. and S. Lemeshow (2000). Applied logistic regression (Wiley Series in probability and statistics) (2 ed.). Wiley-Interscience Publication.

Illian, J. B., S. Martino, S. Sørbye, J. B. Gallego-Fernández, M. Zunzunegui, M. P. Esquivias, and J. M. J. Travis (2013). Fitting complex ecological point process models with integrated nested Laplace approximation. Methods in Ecology and Evolution 4, 305–315.

Johnson, D. H. (1980). The comparison of usage and availability measurements for evaluating resource preference. Ecology 61, 65–71.

Jonsen, I., C. McMahon, T. Patterson, M. Auger-Methe, R. Harcourt, M. Hindell, and S. Bestley (2018). Movement behaviour responses to environment: fast inference of individual variation with a mixed effects model. bioRxiv preprint bioRxiv:314690.

Kays, R., M. C. Crofoot, W. Jetz, and M. Wikelski (2015). Terrestrial animal tracking as an eye of life and planet. Science 348, aaa2478.

Kranstauber, B., A. Cameron, R. Weinzerl, T. Fountain, S. Tilak, M. Wikelski, and R. Kays (2011). The movebank data model for animal tracking. Environmental Modelling & Software 26, 834–835.

Leclerc, M., E. Vander Wal, A. Zedrosser, J. E. Swenson, J. Kindberg, and F. Pelletier (2016). Quantifying consistent individual differences in habitat selection. Oecologia 180, 697–705.

Lee, Y. and J. A. Nelder (2004). Conditional and marginal models: another view. Statistical Science 19, 219–238.

Lele, S. R. and J. L. Keim (2006). Weighted distributions and estimation of resource selection probability functions. Ecology 87, 3021–3028.

Lele, S. R., J. L. Keim, and P. Solymos (2017). ResourceSelection: Resource Selection (Probability) Functions for Use-Availability Data. R package version 0.3-2.

Lindgren, F., H. Rue, and J. Lindström (2011). An explicit link between Gaussian fields and Gaussian Markov random fields: The SPDE approach (with discussion). Journal of the Royal Statistical Society. Series B (Statistical Methodology) 73, 423–498.

Lu, Y. and S. L. Zeger (2007). On the equivalence of case-crossover and time series methods in environmental epidemiology. Biostatistics 8, 337–344.

Lunn, D., A. Thomas, N. Best, and D. Spiegelhalter (2000). WinBUGS–A Bayesian modelling framework: concepts, structure, and extensibility. Statistics and Computing 10, 325–337.

Magnusson, A., H. J. Skaug, A. Nielsen, C. W. Berg, K. Kristensen, M. Maechler, K. J. van Bentham, B. M. Bolker, and M. E. Brooks (2017). glmmTMB: Generalized Linear Mixed Models using Template Model Builder.

Manly, B. F., L. L. McDonald, D. L. Thomas, T. L. McDonald, and W. P. Erickson (2002). Resource Selection by Animals: Statistical Design and Analysis for Field Studies (2 ed.). Dordrecht, The Netherlands: Kluwer Academic Publishers.

Matthiopoulos, J., J. Fieberg, G. Aarts, H. L. Beyer, J. M. Morales, and D. T. Haydon (2015). Establishing the link between habitat selection and animal population dynamics. Ecological Monographs 85, 413–436.

Matthiopoulous, J., M. Hebblewhite, G. Aarts, and J. Fieberg (2011). Generalized functional response for species distributions. Ecology 92, 583–589.

McCullagh, P. and J. A. Nelder (1989). Generalized Linear Models. London: Chapman and Hall.

McDonald, T. L., B. F. J. Manly, R. M. Nielson, and L. V. Diller (2006). Discrete-choice modelling in wildlife studies exemplified by northern spotted Owl nighttime habitat selection. The Journal of Wildlife Management 70, 375–383.

Millspaugh, J. J., R. M. Nielson, L. McDonald, J. M. Marzluff, R. A. Gitzen, C. D. Rittenhouse, M. W. Hubbard, and S. L. Sheriff (2006). Analysis of resource selection using utilization distributions. Journal of wildlife management 70, 384–395.

Muff, S., L. Held, and L. F. Keller (2016). Marginal or conditional regression models for correlated non-normal data? Methods in Ecology and Evolution 7, 1514–1524.

Muff, S., A. Riebler, L. Held, H. Rue, and P. Saner (2015). Bayesian analysis of measurement error models using integrated nested Laplace approximations. Journal of the Royal Statistical Society, Applied Statistics Series C 64, 231–252.

Mysterud, A. and R. A. Ims (1998). Functional responses in habitat use: availability influences relative use in trade-off situations. Ecology 79, 1435–1441.

Phillips, S. J., R. P. Anderson, and R. E. Schapire (2006). Maximum entropy modeling of species geographic distributions. Ecological Modelling 190, 231–259.

Plummer, M. (2003). JAGS: A Program for Analysis of Bayesian Graphical Models Using Gibbs Sampling. In Proceedings of the 3rd International Workshop on Distributed Statistical Computing.

R Core Team (2018). R: A Language and Environment for Statistical Computing. Vienna, Austria: R Foundation for Statistical Computing.

Renner, I. W. and D. I. Warton (2013). Equivalence of MAXENT and Poisson point process models for species distribution modeling in ecology. Biometrics 69, 274–281.

Rhodes, J. R., C. A. McAlpine, D. Lunney, and H. P. Possingham (2005). A spatially explicit habitat selection model incorporating home range behavior. Ecology 86, 1199–1205.

Ribeiro Jr, P. J. and P. J. Diggle (2016). geoR: Analysis of Geostatistical Data. R package version 1.7-5.2.

Rue, H., S. Martino, and N. Chopin (2009). Approximate Bayesian inference for latent Gaussian models by using integrated nested Laplace approximations (with discussion). Journal of the Royal Statistical Society. Series B (Methodological) 71, 319–392.

Rue, H., A. Riebler, S. H. Sørbye, J. B. Illian, D. Simpson, and F. Lindgre (2016). Bayesian computing with INLA: A review. ArXiv e-prints. arXiv:1604.00860.

Schielzeth, H. and W. Forstmeier (2009). Conclusions beyond support: Overconfident estimates in mixed models. Behavioral Ecology 20, 416–420.

Signer, J. (2018). amt: Animal Movement Tools. R package version 0.0.4.0.

Signer, J., J. Fieberg, and T. Avgar (2018). Animal movement tools (amt): R package for managing tracking data and conducting habitat selection analyses. Ecology and Evolution. In press.

Simpson, D., H. Rue, A. Riebler, T. G. Martins, and S. H. Sórbye (2017). Penalising model component complexity: A principled, practical approach to constructing priors. Statistical Science 32, 1–28.

Therneau, T. M. (2015a). coxme: Mixed Effects Cox Models. R package version 2.2-5.

Therneau, T. M. (2015b). A Package for Survival Analysis in S. version 2.38.

Thurfjell, H., S. Ciuti, and M. Boyce (2014). Applications of step-selection functions in ecology and conservation. Movement Ecology 2, 4.

Wang, X., Y. Y. Ryan, and J. J. Faraway (2018). Bayesian Regression Modeling with INLA. Boca Raton: Chapman & Hall/CRC.

Warton, D. and G. Aarts (2013). Advancing our thinking in presence-only and used-available analysis. Journal of Animal Ecology 82, 1125–1134.

Warton, D. I. and L. C. Shepherd (2010). Poisson point process models solve the “pseudo-absence problem” for presence-only data in ecology. Annals of Applied Statistics 4, 1383–1402.

Weinberger, I. C., S. Muff, A. Kranz, and F. Bontadina (2016). Flexible habitat selection paves the way for a recovery of otter populations in the European Alps. Biological Conservation 199, 88–95.

Whitehead, J. (1980). Fitting Cox’s regression model to survival data using GLIM. Applied Statistics 29, 268–275.

Wikelski, M. and R. Kays (2018). Movebank: archive, analysis and sharing of animal movement data. Hosted by the Max Planck Institute for Ornithology, https://www.movebank.org.

Yuan, Y., F. Bachl, F. Lindgren, D. Brochers, J. Illian, S. Buckland, H. Rue, and T. Gerrodette (2016). Point process models for spatio-temporal distance sampling data. arXiv preprint arXiv:1604.06013.

